# Combined PARP14 Inhibition and PD-1 Blockade Promotes Cytotoxic T Cell Quiescence and Modulates Macrophage Polarization in Relapsed Melanoma

**DOI:** 10.1101/2024.09.26.615178

**Authors:** Rotem Leshem, Kieran N. Sefton, Chun Wai Wong, I-Hsuan Lin, Dervla T. Isaac, Mario Niepel, Adam F.L. Hurlstone

## Abstract

Programmed Cell Death 1 (PD-1) signaling blockade effectively restores immune surveillance to treat melanoma. However, chronic interferon-gamma (IFNγ) -driven immune homeostatic mechanisms in melanoma cells contribute to immune evasion and acquired resistance. We previously demonstrated that poly ADP ribosyl polymerase 14 (PARP14), an IFNγ-responsive gene product, in part mediates IFNγ-driven resistance, as its inhibition prolonged PD-1 blockade responses in preclinical models. Nevertheless, PARP14 inhibition alone could not achieve full tumor clearance, indicating additional resistance mechanisms. Herein, we identify a robust PARP14 catalytic inhibitor (PARP14i) gene signature associated with improved patient survival. We elucidate immune and tumor cell adaptations to PARP14 inhibition combined with PD-1 blockade using preclinical models. This combination therapy suppressed tumor-associated macrophages while increasing pro-inflammatory memory macrophages. Notably, it preserved cytotoxic T cell function by inducing a quiescent state, mitigating terminal exhaustion. Despite enhancing immune responses, adaptive resistance mechanisms emerged in tumor cells, engaging alternative immune evasion pathways. These findings indicate that this combination therapy primes immune cells for further therapeutic intervention, providing a promising strategy to overcome resistance and optimize treatment outcomes.

**Summary:** While immune checkpoint inhibitors, such as α-PD-1 therapy, provide significant clinical benefits, many tumors develop resistance and relapse. Re-treatment with α-PD-1 often fails because T cells lose their function (a process called exhaustion), and immune-suppressing cells like tumor-associated macrophages (TAMs) increase in the tumor environment, enabling tumors to evade immune attack.

Recent studies have shown that a small molecule inhibitor of PARP14 can restore sensitivity to α-PD-1 therapy by remodeling the tumor environment. This includes increasing immune cell infiltration and improving antigen presentation, which helps the immune system recognize tumor cells.

Using single-cell RNA sequencing, we discovered that this therapy induces a new state of T cell dormancy, where cytotoxic T cells temporarily stop attacking but retain the potential to be reactivated by additional treatments. This finding opens new opportunities to sustain immune responses and improve outcomes for patients who relapse after immunotherapy.

**Graphical Summary:** 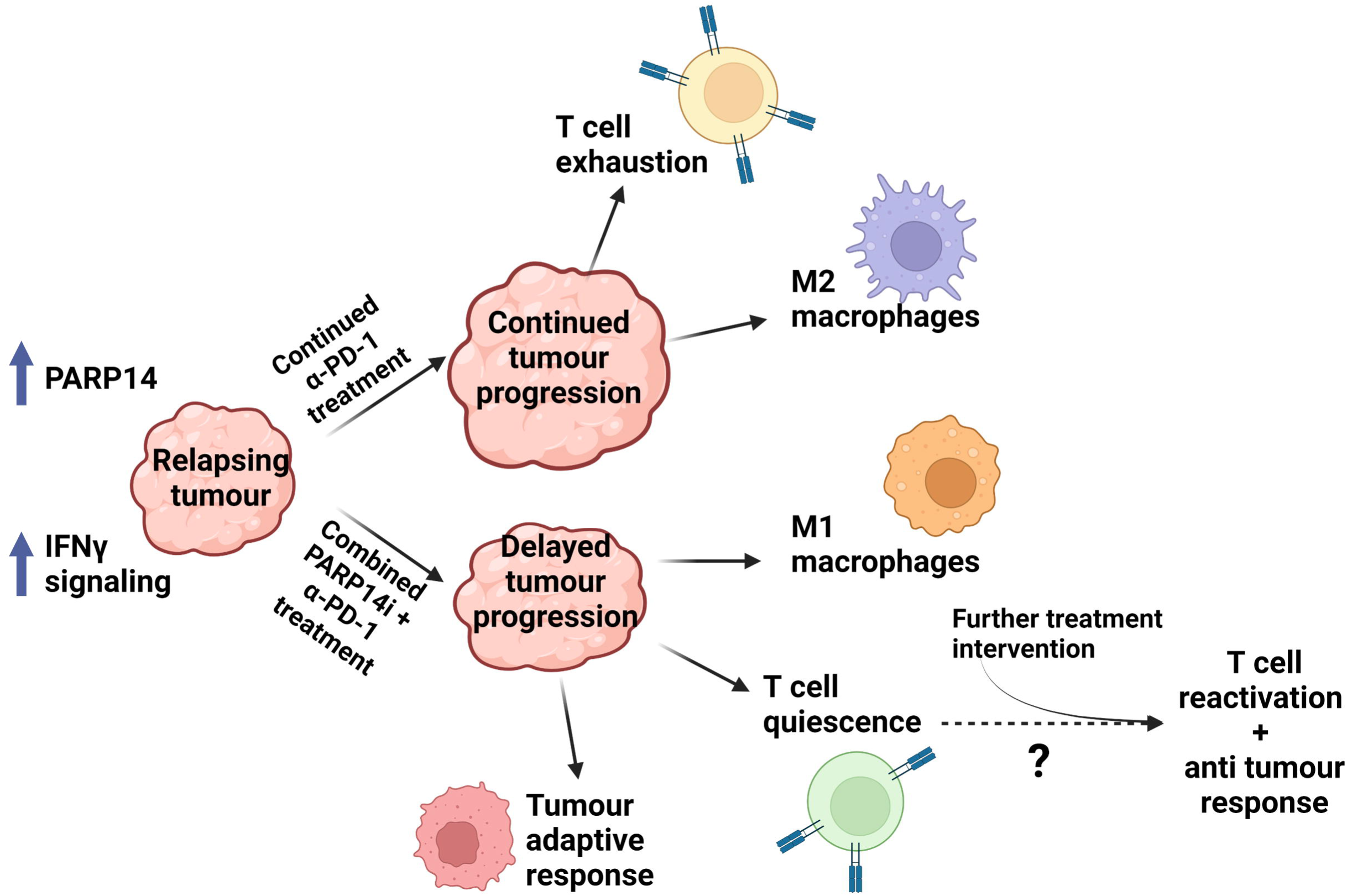

## Introduction

Drugs antagonizing the PD-1 signaling axis by disrupting the interaction between PD-1 on exhausted T cells and Programmed Cell Death Ligand 1 (PD-L1) on tumor and stromal cells have vastly improved outcomes for high-risk and late-stage melanoma patients. However, among responders to PD-1 blockade therapy, approximately 25% will relapse within two years of initiating treatment [1]. Little can be done at present for these patients. Analysis of relapsing tumors has revealed that 30% are characterized by intrinsic—that is, ligand-independent— interferon-gamma (IFNγ) signaling [2]. Intrinsic IFNγ signaling in cancer cells has been shown to follow chronic IFNγ stimulation and to entail epigenetic remodeling wherein signal transducer and activator of transcription 1 (STAT1) and Interferon Regulatory Factor 3 (IRF3) permanently occupy the regulatory sites of a subset of interferon-stimulated genes (ISG), including notably those promoting T cell dysfunction [3]. Indeed, the ISG selectively upregulated in chronically stimulated cancer cells (ISG resistance signature or ISG.RS) predicts poor survival in patients and poor response to PD-1 blockade [4].

Our previous work revealed a contribution from PARP14 in mediating chronic IFNγ stimulation-driven PD-1 blockade resistance in preclinical models [5]. One model we developed entailed implantation of the mouse rapidly accelerated fibrosarcoma homolog B (BRAF) and Phosphatase and Tensin homolog (PTEN) mutant melanoma cell line YUMM2.1 [6] into the hypodermis of syngeneic C57/BL6 mice, and then treating these mice with a short course of anti-PD-1 (α-PD-1) antibodies when tumors became palpable. In tumors that resumed growth following α-PD-1 treatment, we observed increased expression of multiple ISG, including ISG.RS genes, relative to control (untreated) tumors and to tumors still responding to PD-1 blockade therapy [5]. AI-assisted immune profiling, based on deconvolving gene expression, indicated that relapsing tumors were also relatively more T cell infiltrated, including by T_EM_ and T_CM._ However, these T cells were also more dysfunctional [5]. The regrowth of such tumors was significantly delayed—but not abolished— and survival lengthened by adding a highly potent and selective PARP14 inhibitor RBN012759 (PARP14i, Ribon Therapeutics)[7] to the treatment regimen [5].

This previous study also highlighted that the *PARP14* gene is a STAT1 target and that PARP14 itself is a STAT1 interacting protein [5]. Moreover, in macrophages, PARP14 interaction with STAT1 inhibits the expression of proinflammatory genes [8] but regulates IRF3 recruitment of RNA polII to promoters of IRF3 target genes [9] and is thus positioned to influence the expression of ISG. However, the details are far from clear. Certainly, our previous analysis of gene expression indicated that PARP14 inhibitor treatment of melanoma cells chronically stimulated with IFNγ increased IFN-, TNF-α-, and IL-6-driven inflammatory signaling [5].

Intrigued by the potential of PARP14 to influence the expression of ISG in tumor cells but also the failure of PARP14 inhibitor treatment to permanently flatten tumor rebound after α-PD-1 treatment, we undertook a detailed analysis of tumor composition and gene expression changes attending PARP14 inhibitor treatment to identify other targets for intervention which we now present below.

## Materials and Methods

### Reagents

Recombinant human IFNγ (PHC4031, Gibco) and recombinant mouse IFNγ (PMC4031, Gibco). PARP14 inhibitor RBN012759 [7] and PARP14 PROTAC RBN012811 (Ribon Therapeutics) [10].

### Cell lines

Human melanoma cell lines A375 and 501-Mel (provided by Claudia Wellbrock, The University of Manchester), MC38 cells (provided by Santiago Zelenay, The University of Manchester) and YUMM2.1 cells (provided by Richard Marais, The University of Manchester) were maintained in RPMI-1640 (Sigma–Aldrich) supplemented with 10% v/v fetal bovine serum (Life Technologies) and 1% w/v penicillin-streptomycin (Sigma–Aldrich), at 37 °C in a 5% v/v CO_2_ humidified incubator. Cell lines authenticated by STR profiling; cultures routinely tested for mycoplasma. Cells were treated for two weeks with 50 IU/mL IFNγ, refreshed every 2-3 days, and subsequently treated with 500nM DMSO (Sigma–Aldrich); 500nM PARP14i; or 100nM PARP14 PROTAC for 48h.

### Mouse tumor implants

Mice were housed in the Biological Services Facility of The University of Manchester on a 12/12 h light/dark cycle and given ad libitum food (Bekay, B&K Universal, Hull, UK) and water. All procedures were approved by the University of Manchester’s animal welfare ethical review board and performed under relevant Home Office licenses according to the UK Animals (Scientific Procedures) Act, 1986.

8–12-week-old C57BL/6 female mice (ENVIGO) were allowed 1-week to acclimatize. YUMM2.1 cells (5 × 10^6^ cells) in 100 μL serum-free RPMI-1640 were subcutaneously injected into the left flank. Tumor volumes (height x width x length caliper measurements) and mouse weights were monitored every 2–3 days. When tumors reached a volume of 80 mm^3^, mice were administered two doses of 300 μg α-PD-1 antibody (BioXCell) in 100 μL InVivoPure pH 7.0 Dilution Buffer (BioXCell) via intraperitoneal (i.p.) injection administered at 3-day intervals. Mice were culled if tumors remained under 100 mm^3^ (total responders) or conversely reached 800 mm3 (non-responders). Otherwise, mice with tumors recovering to 140 mm^3^ were randomized to a second round of treatments (re-treatment): 2 doses of 300 μg of rat isotype control antibody IgG2b (BioXCell) (IgG group) or α-PD-1 antibody (BioXCell) (α-PD-1 group) in 100 μL InVivoPure pH 7.0 Dilution Buffer (BioXCell) via i.p. injection at three-day intervals, 14 doses of 500 mg/Kg of PARP14I by oral gavage twice a day (PARP14i group), or a combination of α-PD-1 i.p. and PARP14i gavage in the same doses (Combination group). PARP14i was dissolved in 0.5% w/v methylcellulose (Sigma–Aldrich) + 0.2% v/v Tween 80 (Sigma–Aldrich). Dose volume - 10 mL/kg.

### Tumor immune infiltrate analysis by flow cytometry

Tumors were dissected upon reaching 250-400mm^3^ following retreatment and incubated for 45 minutes with 100 μg/mL Liberase (Sigma–Aldrich) in serum-free media at 37 °C, then pushed through a 100 μM nylon cell strainer. The cell suspension was stained for 20 min with Zombie UV™ Fixable Viability Kit (BioLegend) in PBS. Subsequently, Fc receptors were blocked, and cells were stained with a surface stain antibody mix. Cells were fixed and permeabilized using the Foxp3/Transcription Factor Staining Buffer Set (eBioscience) following manufacturer instructions. Samples were measured on a BD Fortessa flow cytometer (BDBiosciences), and data was collected using BD FACSDiva™ software. For all antibodies, a non-stained cell sample and appropriate fluorescence minus one control were analyzed as well. Data were analyzed using FlowJo version 8.7.

Antibodies: CD16/32 (clone 93), CD45 (Cat. 103132), CD3ε (Cat. 100306), CD4 (Cat. 100552), CD8α (Cat. 100742), PD-1 (Cat. 135231), LAG-3 (Cat. 125224), TIM-3 (Cat. 119721), Ki-67 (Cat. 652420), Granzyme B (Cat. 372214), CD69 (Cat. 104531), and CTLA-4 (Cat. 106338) from Biolegend. Tox (Cat. 12-6502-82) was purchased from eBioscience. Gating strategy as described in SUPPLEMENTARY FIGURE 5.

### Single-cell RNAseq sequencing

#### Tumor processing

Tumors were collected on day 7 of retreatment into serum-free RPMI on ice, then minced into 1mm pieces and incubated at 37 °C up to 60 min in Liberase TL (Millipore Sigma) with rocking according to manufacturer instructions. Digestion was stopped using fetal bovine serum, and tissue was further dissociated using a wide-bore pipette and filtered through a cell strainer to achieve a single-cell solution. Immune population enrichment was done by CD45 mouse Microbeads (Miltenyi) according to manufacturer instructions. Following sorting, both populations were counted, and ∼5% of CD45 negative cells were spiked back into the CD45 positive fraction. Cells were fixed for 18 hours, then quenched and stored at - 80°C in Enhancer according to 10x Genomics Tissue Fixation & Dissociation for Chromium Fixed RNA Profiling protocol (CG000553 Rev A) until all samples were collected.

#### Single-cell isolation and library construction

Gene expression libraries were prepared from formaldehyde-fixed samples using the Chromium X and Chromium Fixed RNA kit, Mouse Transcriptome, 4 rxns x 16 BC (10x Genomics, Inc.) according to the manufacturer’s protocol (CG000527 Rev C). Illumina-compatible sequencing libraries were constructed by adding P5, P7, i5, and i7 sample indexes and Illumina Small Read 2 sequences to the 10x barcoded, ligated probe products via Sample Index PCR followed by a SPRIselect size-selection.

#### Sequencing

The resulting sequencing library comprised standard Illumina paired-end constructs flanked with P5 and P7 sequences. The 16 bp 10x Barcode and 12 bp UMI were encoded in Read 1, while Read 2 sequenced the ligated probe insert, constant sequence, and the 8bp Probe Barcode. Sample indexes were incorporated as the i5 and i7 index reads. Paired-end sequencing (28:90) was performed on the Illumina NovaSeq6000 platform.

### Data Analysis

#### Raw data processing and cell filtering

Raw sequencing data were processed using the 10x Genomics Cell Ranger pipeline (v7.0.0). Base call (BCL) files generated by the sequencer were demultiplexed and converted to FASTQ files using “cellranger mkfastq”. The FASTQ files were then mapped against the pre-built Mouse reference package from 10X Genomics (mm10-2020-A) using “cellranger multi” to demultiplex and produce the gene-cell barcode matr-ix for individual samples. The single-cell data were processed in R environment (v4.1) following the workflow documented in Orchestrating Single-Cell Analysis with Bioconductor [11]. Briefly, for each sample, the HDF5 file generated by Cell Ranger was imported into R to create a SingleCellExperiment object. A combination of median absolute deviation (MAD), as implemented by the “isOutlier” function in the scuttle R package (v1.4.0), and exact thresholds were used to identify and subsequently remove low-quality cells before data integration.

#### Data integration, visualization, and cell clustering

The log-normalized expression values of the combined data were re-computed using the “multiBatchNorm” function from the bachelor R package (v1.10.0). The per-gene variance of the log-expression profile was modeled using the “modelGeneVarByPoisson” function from the scran R package (v1.22.1), and the top 5000 highly variable genes were selected. The mutual nearest neighbors (MNN) approach implemented by the “fastMNN” function from the bachelor R package was used to perform batch correction. The first 50 dimensions of the MNN low-dimensional corrected coordinates for all cells were used as input to produce the t-stochastic neighbor embedding (t-SNE) projection and uniform manifold approximation and projection (UMAP) using the “runTSNE” and “runUMAP” functions from the scater R package (v1.22.0) respectively. Putative cell clusters were identified using the Leiden algorithm from the igraph R package (v1.3.0).

#### Marker genes and differential expression analysis

Cluster-specific markers were identified using the “findMarkers” function from the scran R package, which performs pairwise t-tests between clusters. Differential expression analysis between treatment and control (IgG) was performed on pseudo-bulk samples using the quasi-likelihood pipeline from the edgeR R package (v3.36.0). Genes with a false discovery rate (FDR) below 5% were considered differentially expressed.

#### Pathway analysis

Pathway enrichment analysis of differentially expressed genes was performed using the R package enrichR (v3.0). Pathway results were ranked by the combined scores (calculated as log(P-value) multiplied by z-score), and pathways with an adjusted P-value < 0.05 were considered significant. Further gene set enrichment analysis GSEA was done using Metascape web tool [12] and plotted as -log_10_P.

#### Subclustering and further analysis

Cells were divided into functional cell type groups and further subclustered using the Louvain method for community detection [13]. Trajectory inference was made using the Slingshot R package [14], and the association of our PARP14i signature with survival in immune checkpoint blockade therapy(ICBT) cohorts, as well as a comparison of high and low-expressed genes in specific clusters, was assessed using Tumor Immune Dysfunction and Exclusion (TIDE) web tool [15]. Sample RL1i projection on UMAP was done using the ProjecTILs R package [16].

#### Data Availability

Single-cell RNAseq data have been deposited in the ArrayExpress database at EMBL-EBI under accession number E-MTAB-14471. Bulk RNA sequencing was performed as previously described [5], with data available from the ArrayExpress database under accession numbers <u>E-</u> <u>MTAB-12195</u> and <u>E-MTAB-12872</u>.

## Results

### A genetic signature reflecting the modulating effect of PARP14i on chronic IFNγ stimulation predicts response to α-PD-1 in clinical cohorts

We and others have previously implicated PARP14 in modulating type II interferon signaling [5,8,17]. Following chronic stimulation of several mouse and human cancer cell lines with IFNγ and then subsequent treatment for 48 hours with a PARP14 catalytic inhibitor (PARP14i), PARP14 proteolysis targeting chimera (PROTAC), or aqueous Dimethyl sulfoxide (DMSO; drug solvent control), we assessed differences in gene expression by bulk RNA sequencing (RNAseq). Comparison of the top 50 differentially expressed genes within each cell model revealed a shared expression pattern, which we narrowed down to a PARP14i signature consisting of 30 genes (**FIGURE 1A, <u>SUPPLEMENTARY FIGURE 1A and SUPPLEMENTARY TABLE 1</u>**). Pathway enrichment analysis performed by Metascape [12] and ARCHS4 [18] revealed that among this signature were genes strongly correlated with response to IFNγ signaling, with an emphasis on STAT/IRF function (**FIGURE 1B <u>and SUPPLEMENTARY FIGURE 1B</u>**). However, we found this signature is distinct from the genes contained within the IFNγ signaling GO term, with less than half of the PARP14i signature displaying overlap. We then investigated whether this PARP14i signature could predict response to α-PD-1 therapy by employing Tumor Immune Dysfunction and Exclusion (TIDE) analysis [15]. High expression of genes within this signature was associated with substantial improvement in overall patient survival across several clinical cohorts receiving immune checkpoint blockade therapy [19–21] (**FIGURE 1C**), underscoring the prognostic value of this signature and supporting the potential therapeutic benefit of PARP14 inhibition.

**Figure 1.**
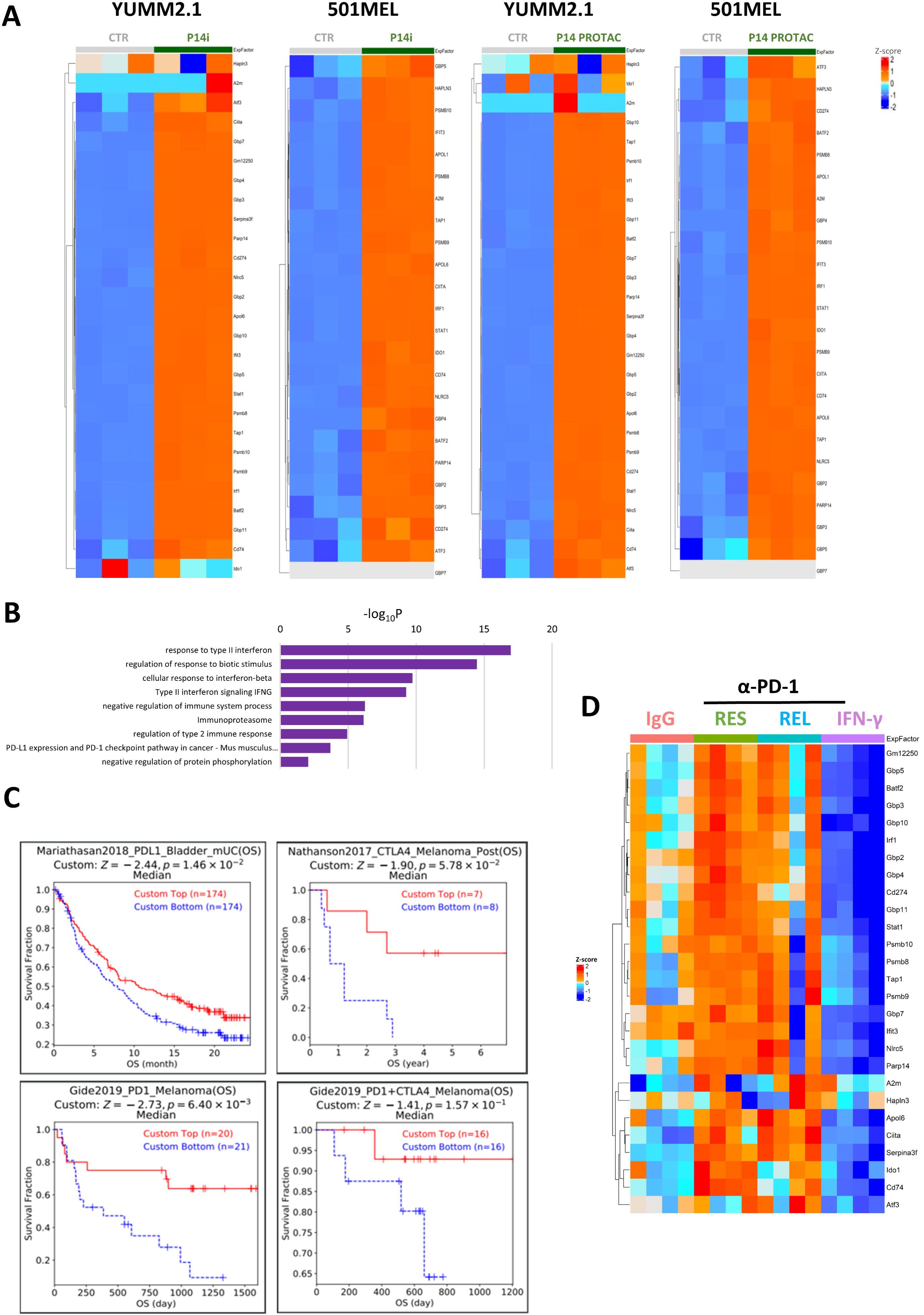
Identification of a genetic signature associated with PARP14 inhibition. YUMM2.1 and 501MEL cells treated for 48h *in vitro* with either DMSO, 100nM PARP14 inhibitor (RBN012759), or 500nM PARP14 PROTAC (RBN012811); *n* = 3. **A.** Bulk RNAseq analysis of a PARP14 inhibition signature identified across human and mouse cell lines. Heatmap of treatment vs control per cell line, colors representing row-scaled z-score expression high (red) to low (blue). **B.** Gene set enrichment analysis (GSEA) of 30 PARP14i signature genes identified by comparison of bulk RNAseq from human and mouse cell lines, chronically treated with IFN-γ followed by 48 hours of PARP14 inhibition (Metascape analysis, values presented as -log_10_ of P value). **C.** Kaplan-Meier (KM) curves showing cancer patients with high Tumor Immune Dysfunction and Exclusion (TIDE) scores of PARP14i signature genes having better overall survival (OS) rates in representative studies. Red indicates patients with high (top 50%) TIDE scores and blue indicates low (bottom 50%) TIDE scores. **D.** Bulk RNAseq analysis on naïve YUMM2.1 tumors treated with IgG or α-PD-1. α-PD-1-treated cohort was split into relapsed (REL) and responders (RES). Compared to YUMM2.1 tumors pre-treated with chronic IFN-γ stimulation.

Furthermore, we performed RNAseq analysis on subcutaneous tumors comprising YUMM2.1 cells dissected from mice treated with either IgG or α-PD-1, dividing the latter into those either relapsing or responding to the treatment. Although the PARP14i signature was upregulated in α-PD-1-treated samples, it did not distinguish between treatment response and relapse (**FIGURE 1D**). For further comparison, we also analyzed IgG-treated tumors arising from YUMM2.1 cells pre-treated *ex vivo* with chronic IFNγ before implantation and found that the PARP14i signature was significantly downregulated (**FIGURE 1D**), consistent with a tumor microenvironment (TME) that is T cell cold and inherently resistant to α-PD-1 [5].

### Analysis of changes to the cellular composition of YUMM2.1 tumors relapsing following α-PD-1 therapy driven by subsequent PARP14 inhibitor treatment

To investigate further the ability of PARP14i to improve α-PD-1 response, we turned to our previously described mouse model of acquired resistance [5]: YUMM2.1 cells were implanted subcutaneously, and upon tumors reaching ∼80mm^3^ in volume, two doses of α-PD-1 antibodies were given three days apart (**FIGURE 2A <u>and SUPPLEMENTARY FIGURE 2</u>**.**A)** Tumor volumes were routinely recorded (**<u>SUPPLEMENTARY TABLE 2</u>**), and at ∼100-150mm^3^, animals were randomly allocated into the following re-treatment groups: IgG2a (n=4); α-PD-1 (n=3); PARP14i alone (n=3); and a combination of α-PD-1 + PARP14i (n=4). Animals were sacrificed following seven days of re-treatment, and tumors were subsequently processed for single-cell RNA sequencing (scRNAseq). Following tumor dissociation and enrichment for CD45^+^ cells, 5% from the CD45^−^ cell fraction was added to ensure tumor cell representation in our analyses. Cells were fixed and frozen until all tumor samples were collected; then, samples were barcoded, multiplexed, and libraries prepared and sequenced.

**Figure 2.**
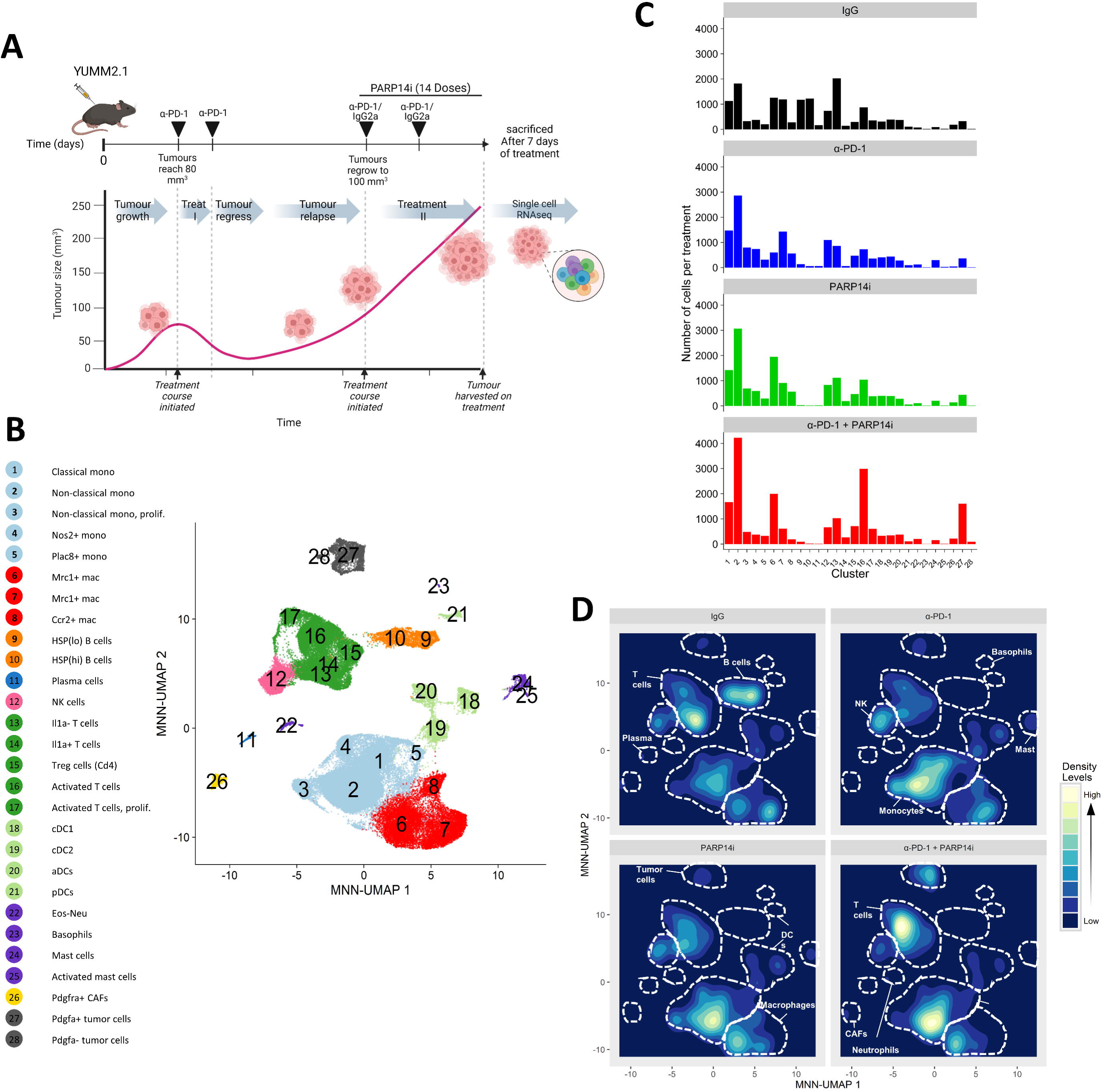
Analysis of the cellular composition of YUMM2.1 tumors relapsing following α-PD-1 antibodies with and without PARP14 inhibitor treatment. **A.** YUMM2.1 cells were implanted subcutaneously into the flank of female C57BL/6 mice. Once tumors reached ∼80mm^3^ in volume, treatment of α-PD-1 antibodies was commenced for a total of two doses, 3 days apart. Tumor shrinkage and regrowth was monitored. When tumors reached a volume between ∼100-150mm^3^, a second round of treatment commenced, lasting 7 days. Tumors were subsequently processed for single-cell RNAseq. Graphical summary of experimental timeline, showing tumor inoculation and treatment regime (top), and illustration of tumor growth (bottom). Created in BioRender. Leshem, R. (2023) BioRender.com/r18g555. **B.** Mutual nearest neighbor (MNN) corrected Uniform Manifold Approximation and Projection (UMAP) of 13 tumor samples (*n*=65,476 cells). Clustered using Leiden unsupervised clustering and colored by cell types. Cluster identifies below. **C.** Cell number changes in each cluster by treatment: three mice IgG2a; three mice α-PD-1; three mice PARP14i (RBN012759) alone; and four mice a combination of α-PD-1 + PARP14i (RBN012759). **D.** Cell density projections by treatment.

In total, fourteen samples of tumor immune cells were sequenced. Initial analysis of sequencing data identified one of the IgG samples (RL11i) to be anomalous, which upon further review of its corresponding tumor growth curve was attributed to it being, in reality, a non-responder to the initial α-PD-1 treatment, and therefore, it was excluded from subsequent analyses (**<u>SUPPLEMENTARY FIGURE 2A-C</u>**). Leiden unsupervised clustering across the remaining thirteen treated samples identified twenty-eight clusters based upon gene expression patterns, with a diverse range of cell types represented across samples and treatment groups (**FIGURE 2B, <u>SUPPLEMENTARY FIGURES 2C and D and SUPPLEMENTARY TABLES 3-4</u>**). The following cell types were identified: Monocytes (Clusters 1-5); Macrophages (6-8); B cells (9-10); Plasma cells (11); Natural Killer (NK) cells (12); T cells (13-17); Dendritic cells (18-21); Granulocytes (22-25); Cancer-Associated Fibroblasts (26); and Tumor cells (27-28). To characterize the individual clusters within these groups, we manually validated cell type identity by comparing the expression of known markers across clusters (**<u>SUPPLEMENTARY TABLES 5-8</u>**).

Macrophages, Monocytes, and T cells were the most abundant leukocyte populations across all treatment groups, with distinct subpopulations displaying treatment-specific enrichment (**<u>SUPPLEMENTARY FIGURE 2E</u>**). Notably, T cells were particularly abundant in combination-treated samples, with increased prevalence of clusters 14 and 16 and depletion of cluster 13, which was highly enriched in IgG-treated samples (**FIGURE 2C**). Similarly, analysis by cell density plots revealed a shift from cluster 7 in macrophages, prominent in α-PD-1-treated samples, towards cluster 6 in the PARP14i and combination treatments (**FIGURE 2D**). Furthermore, we observed IgG-specific enrichment and depletion of B cells and monocytes, respectively. Similarly, across our treatment cohort, few neutrophils and eosinophils were identified, and these could not be well separated, yet we found the sequenced tumor sample, which did not respond to α-PD-1 to be highly enriched in neutrophils (**<u>SUPPLEMENTARY FIGURE 2B</u>**), consistent with a documented role of neutrophils in promoting rapid tumor growth [22].

### PARP14 inhibitor treatment increases M1 macrophage polarization to generate a memory-like state

Given the previous demonstration that PARP14i can suppress the reprogramming of Tumor-Associated Macrophages (TAMs) [23] that are widely considered to be pro-tumorigenic [24] and mimics the effects of α-PD-1 treatment on macrophages in renal tumor explants [7], we interrogated the effects of PARP14i upon the macrophage populations in the TME within our specimens. By further subclustering clusters 1-8 from the initial analysis, we created a new MonoMac UMAP representation detailing a clear division of these subclusters into those that exhibited characteristics of classically activated (M1) macrophages (Left) or alternatively activated (M2) macrophages (Right) (**FIGURE 3A** <u>and **SUPPLEMENTARY TABLES 6, 11**</u>).

**Figure 3.**
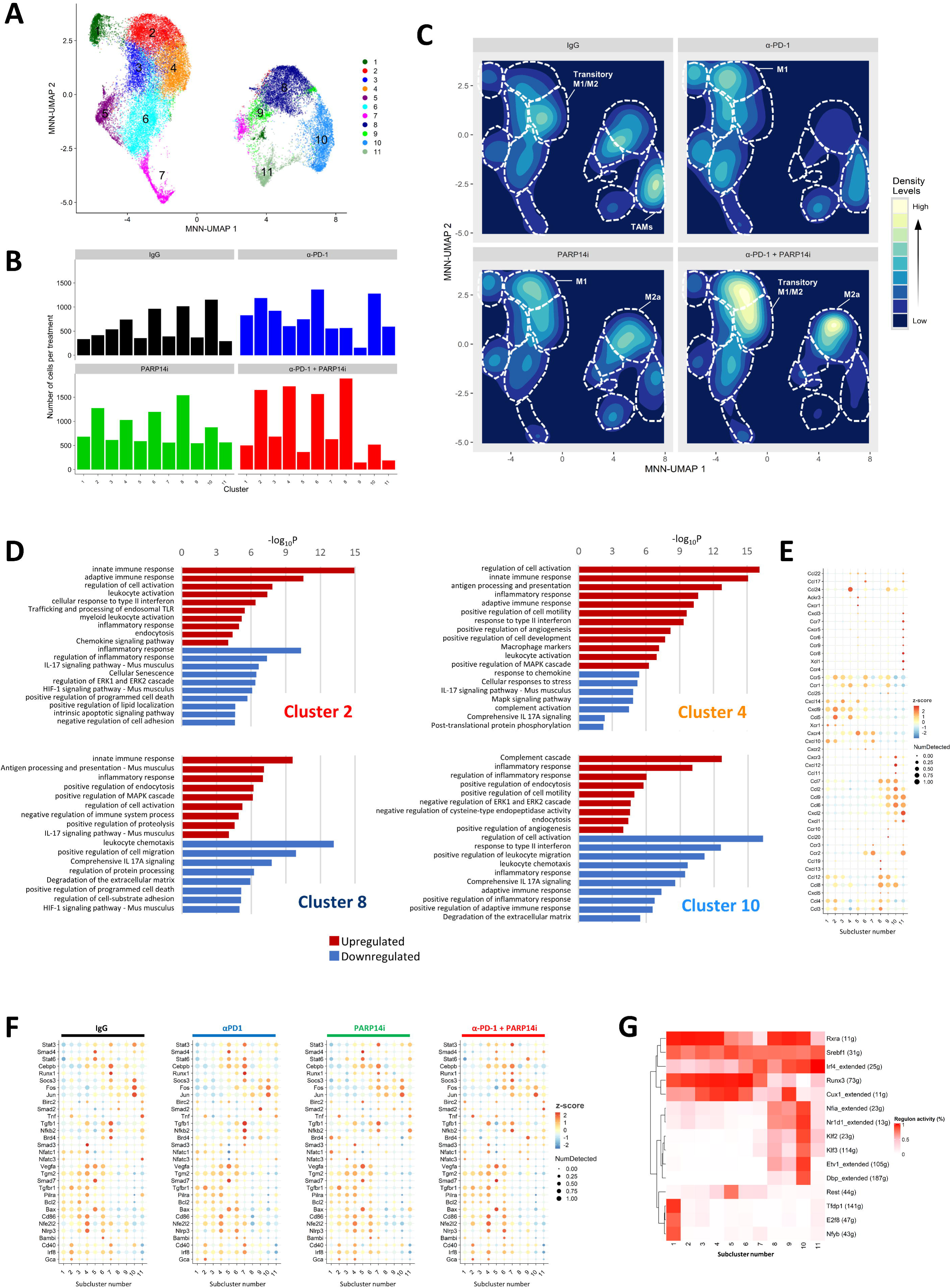
PARP14 inhibitor treatment increases M1 macrophage polarization to generate a memory-like state. **A.** MNN corrected UMAP of monocytes and macrophages (clusters 1-8 of main data, *n*=34,705). Subclustered using Louvain method of community ordering and colored by subclusters. **B.** Cell number changes in each subcluster by treatment: IgG2a; α-PD-1; PARP14i (RBN012759); combination treatment α-PD-1 + PARP14i (RBN012759). **C.** Cell density projections of both monocytes and macrophages (colloquially referred to as MonoMacs) by treatment. **D.** Macrophage subcluster identity-subclusters 2, 4, 8 and 10. GSEA of up-(FDR=0.05, logFC > 1, marked in red) and downregulated (FDR=0.05, logFC < −1, marked in blue) genes (Metascape analysis, values presented as -log_10_ of P value). **E.** Cytokine and Chemokine gene expression presented as high (red) to low (blue) average expression and ratio of cells expressing (circle size), across Macrophage subclusters. **F.** Gene expression presented as high (red) to low (blue) average expression and ratio of cells expressing (circle size), across Macrophage subclusters in different treatments. **G.** Binarized heatmap of 10% random subsampling of each MonoMac subcluster by relative regulon activity (SCENIC analysis).

Analysis of cell type proportions within this subclustering revealed treatment-specific changes. Combination treatment notably shifted macrophage polarization from M2 towards M1 (**FIGURE 3B**). Further inspection of gene enrichment analysis of Differentially Expressed Genes (DEGs), applied alongside cell density mapping, allowed us to robustly identify subclusters most affected by treatment (**FIGURE 3C & D**). This demonstrated that subcluster 10— the most abundant cluster in the IgG-treated samples— is effectively depleted by Combination treatment. This subcluster consists primarily of TAMs, which are known to facilitate tumor progression and metastasis, and as such, their reduction by Combination treatment disrupts pro-tumoral mechanisms.

These anti-tumoral effects are further supported by the expansion of specific macrophage subsets. We observed an increase in M1 macrophages (subcluster 2), signifying an enhanced pro-inflammatory response that bolsters anti-tumor immunity. This increase highlights the ability of Combination treatment to promote a macrophage phenotype that is associated with effective immune surveillance and cytotoxic activity. Similarly, the increased abundance of a transitory M1/M2 macrophage state (subcluster 4) further illustrates the dynamic changes in macrophage populations, highlighting cellular plasticity modulated by Combination treatment to influence the balance between inflammatory and immunoregulatory activities. Concurrently, macrophages in subcluster 8, most closely associated with an M2a macrophage identity, play a nuanced role in the TME under combination treatment. These macrophages exhibit high expression of *Cxcl13* (**FIGURE 3E**), which can mediate both pro- and anti-tumoral effects. CXCL13 interaction with CXCR5 can recruit cytotoxic T cells to the TME, promoting tumoricidal factor secretion and supporting the formation of tertiary lymphoid structures [25,26]. Conversely, this axis can also enhance IL-10 secretion by tumor cells, suppressing the pro-inflammatory functions of antigen-presenting cells (APCs) or recruiting immunosuppressive cells [27]. In parallel, the expression of *Ccl7* and *Cxcl1* by M2a may also contribute to pro-tumorigenic effects, such as promoting angiogenesis and invasion [28,29], reflecting the dual functionality of this macrophage subset in the TME.

The overall increase in M1 macrophages following Combination treatment is linked to elevated expression of pro-inflammatory genes (e.g., *Tnf* and *Stat3*) (**<u>FIGURE 3F</u>**), particularly in subclusters 2 and 4, while the decreased expression of *Stat6* and *Jun* correlates with their reduced transcriptional activity and cellular activation. These changes suggest subclusters 2 and 4 may represent memory-like cells with the potential for re-activation [30]. In contrast, PARP14i treatment results in a marked downregulation of *Nfatc1*, *Nfatc3,* and *Smad4* across subclusters, reducing *Tgfb1* signaling in M2 macrophage clusters, as evidenced by lower *Sox2* and *Sox4* expression (**FIGURE 3F**) [31]. The depletion of key drivers of M2 macrophage polarization such as *Tgm2*, *Vegfa,* and *Cebpb* [32] in clusters 8-11 by Combination treatment aligns with an overall reduction in M2 macrophage populations. In the context of subcluster 8, however, which is enriched by Combination therapy, the reduced levels of anti-inflammatory gene expression by these cells (**FIGURE 3F**) posits them as a transitional macrophage state with reduced features of classical M2 macrophages, reinforcing a Combination treatment-driven transition towards a more hostile tumor environment.

To further explore the molecular underpinnings of macrophage polarization, we applied <u>S</u>ingle-<u>C</u>ell r<u>E</u>gulatory <u>N</u>etwork Inference and <u>C</u>lustering (SCENIC) analysis [33] to evaluate transcriptional regulatory programs within these subclusters (**FIGURE 3G**). Here, we identified regulons that effectively delineated M1 (*Runx3* and *Cux1*) and M2 (*Irf4*) macrophages. Additionally, six regulons associated with M2 macrophages (*Nfia, Nr1d1, Klf2, Klf3, Etv1, Dbp*) were most strongly associated with subcluster 10. In the context of Combination treatment, the depletion of subcluster 10 resulted in a significant reduction in the activity of these regulons. However, this reduction may be offset by the enrichment of subcluster 8, indicative of a directional shift in this treatment towards an M1-like phenotype. These findings suggest that the overall balance between M1 and M2 macrophages is shifting towards a pro-inflammatory, anti-tumor state in Combination treatment. Notably, subclusters 2 and 4, despite their relatively low regulon activity beyond the M1-defining features, may represent more memory-like macrophage states.

### PARP14 inhibitor treatment in combination with α-PD-1 therapy drives activated cytotoxic T cells and limits their transition to dysfunctional states

PARP14 (also known as ARTD8) has been shown to interact with STAT6, a transcription factor activated by interleukin-4 (IL-4), to promote Th2 cell differentiation [34], while Hoon Cho and colleagues demonstrated that PARP14 is involved in the regulation of antibody responses, particularly IgA production, through promoting Th17 differentiation [35]. Both Th2 and Th17 cells are implicated in immune dysfunction in tumors [36] Moreover, our previous work demonstrated that chronic PARP14i treatment of stimulated T cells resulted in increased percentages of IFNγ- and TNFα-producing CD4 and CD8 T cells, accompanied by decreases in TGFβ and IL-10 [5], consistent with a shift toward a type 1 pro-inflammatory T cell response in the TME.

To further characterize specific T cell identities and assess their functionality, we conducted subclustering of clusters 12-17 from the initial clustering (**FIGURE 4A**). By combining NK and T cells for further subclustering, we can be confident that our new T cell clusters are free from any contaminating readouts due to the presence of NK T cells. This subclustering can be split broadly into NK and NKT cells (1-5) and T cells (6-19), with key subpopulations including naïve T cells (7 and 19), regulatory T cells (Tregs) (10 and 11), and cytotoxic T cells (13-18). Comparing the proportions of each T cell subcluster revealed treatment-specific changes in cellular identity and function (**FIGURE 4B <u>and SUPPLEMENTARY TABLE 9</u>**).

**Figure 4.**
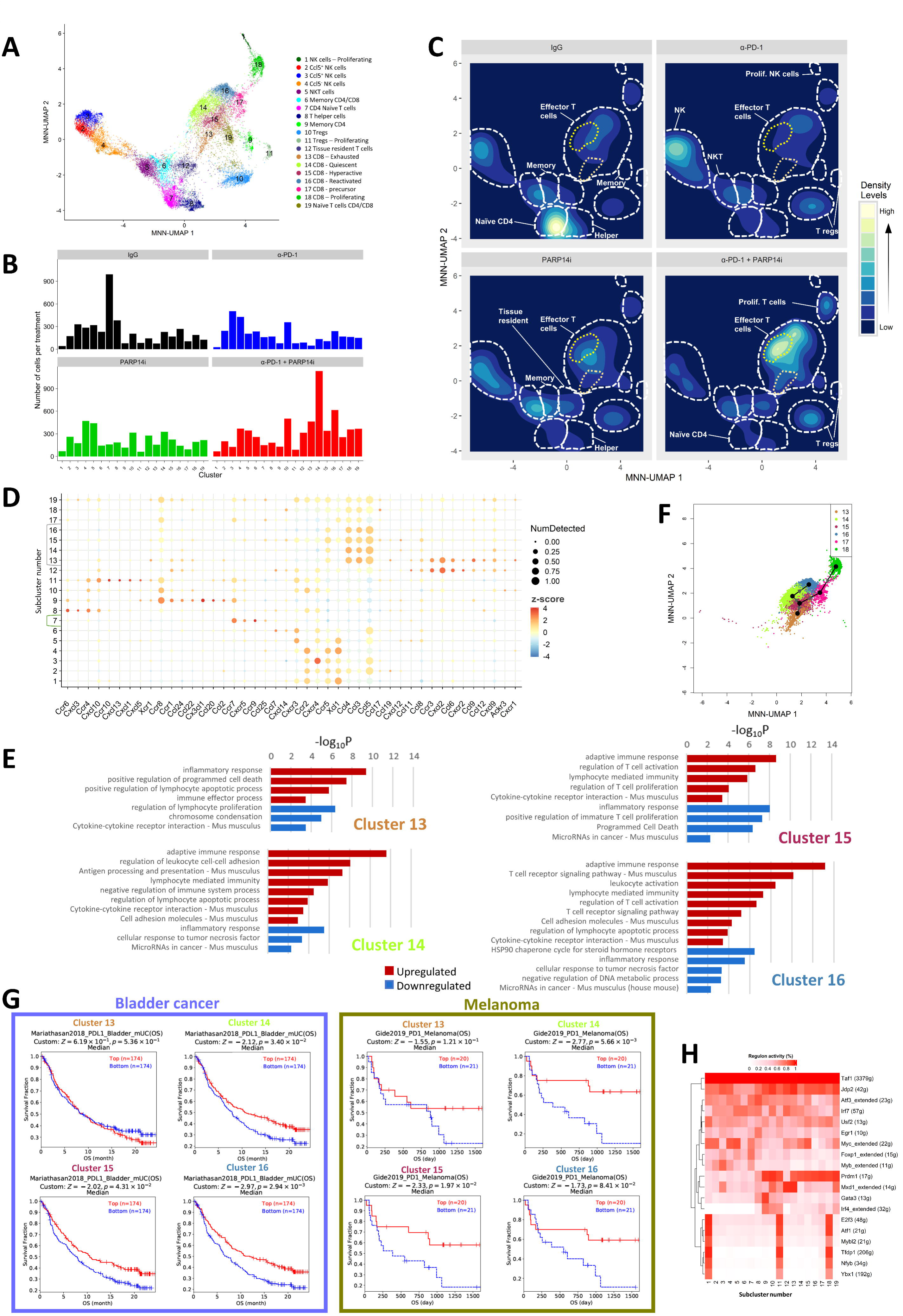
PARP14 inhibitor treatment in combination with α-PD-1 therapy drives activated cytotoxic T cells but limits transition to dysfunctional states. **A.** MNN corrected UMAP of T cell and natural killer (NK) cells (clusters 12-17 main data, *n=18,298*). Subclustered using Louvain method of community ordering and colored by subclusters. **B.** Cell number changes in each T and NK cell subcluster by treatment: IgG2a; α-PD-1; PARP14i (RBN012759); combination treatment α-PD-1 + PARP14i (RBN012759). **C.** Cell density projections of T and NK cells by treatment. **D.** Cytokine and Chemokine gene expression presented as high (red) to low (blue) average expression and ratio of cells expressing (circle size), across T and NK cell subclusters. **E.** T cell subcluster identity-clusters 13-16. GSEA relevant terms of up-(red, FDR=0.05, top 25 genes) and downregulated (blue, FDR=0.05, bottom 25 genes) genes (Metascape analysis, values presented as -log_10_ of P value). **F.** Slingshot trajectory inference analysis of CD8^+^ T cell subclusters 13-18 (*n=6,194*) with trajectory direction indicated by arrows (black). **G.** KM curves showing cancer patients with high TIDE scores of genes from T cell subclusters 13-16 and their effect on OS rates in bladder cancer cohort. Red indicates patients with high (top 50%) TIDE scores and blue indicates low (bottom 50%) TIDE scores. **H.** Binarized heatmap of each T cell subcluster by relative regulon activity (SCENIC analysis).

Cell density mapping (**FIGURE 4C**) highlighted a treatment-dependent decrease in naïve T cells (subcluster 7), which were highly abundant in IgG-treated samples, suggesting their transition into differentiated or activated states upon treatment. In accordance, in IgG-treated tumors, naïve T cells (subcluster 7) exhibited high expression of *Ccr7* and *Ccr9*, which likely mediated their recruitment to the TME (**FIGURE 4D**). CCR7 facilitates migration toward lymphoid tissues via CCL19, a chemokine produced by exhausted CD8+ T cells (subcluster 13) and M2a macrophages (MonoMac subcluster 8) [37]. This interaction may reflect a futile attempt to restock the TME with tumor-inexperienced T cells. However, these naïve T cells exhibited increased *Sell* expression compared to other T cell subclusters, indicative of early activation (**<u>SUPPLEMENTARY FIGURE 3A</u>**) and that IgG-treated tumors suffer from dysfunctional T cell priming by APCs.

Corroborating our hypothesis that the reduction in naïve T cells is due to their differentiation into more active T cell states, we identified a significant enrichment of effector T cells (subclusters 13–16) in the combination treatment group. These cells are recruited and retained in the TME through interactions with pro-inflammatory chemokine ligands, including CCL3, CCL4, and CCL5 (**FIGURE 4D**) [38]. This recruitment is further reinforced by the CXCR5-CXCL13 axis, where CXCL13, predominantly produced by M2a macrophages in our data as previously described (MonoMac subcluster 8), drives cytotoxic T cell recruitment to the TME and aids the formation of tertiary lymphoid structures [25,26]. These structures serve as immune activation hubs, enabling CXCR5-expressing effector T cells to undergo further activation and proliferation, enhancing their cytotoxic function.

Subclusters 13–16 display distinct functional profiles associated with T cell exhaustion, quiescence, hyperactivation, and T cell response, respectively (**FIGURE 4E <u>& SUPPLEMENTARY FIGURES 3B & C</u>**). In the context of anti-tumor activity, T cell subclusters 13-16 are closely interconnected in driving the T cell effector response. Our slingshot trajectory analysis (**FIGURE 4F**) indicates that by entering a state of dormancy (subcluster 14), hyperactive T cells (subcluster 15) sidestep terminal exhaustion (subcluster 13) in the presence of chronic antigen stimulation. Thereafter, quiescent cells can either undergo reactivation (subcluster 16) or persist as a reservoir of memory T cells with retained effector function, poised for re-activation.

TIDE analysis was performed using gene expression signatures arising from each T cell subcluster from subclusters 13-16 (**<u>SUPPLEMENTARY TABLE 10</u>**), revealing that high expression of the gene signature for subcluster 16 (reactivated T cells) was associated with significantly improved survival of patients receiving immune checkpoint blockade therapy for either melanoma or bladder cancer (**FIGURE 4G <u>and SUPPLEMENTARY. FIGURE 3E</u>**) Furthermore, we observed this was also true, albeit to a lesser degree, for the gene signatures arising from subclusters 14 (quiescent T cells) and 15 (hyperactive T cells). On the other hand, the gene signature of T cell subcluster 13 (exhausted T cells) did not associate with positive patient outcomes, in keeping with T cell exhaustion constituting a profound loss of T cell functionality [39]. This suggests that cells within subclusters 14–16 are critical in mounting an effective anti-tumor response through mechanisms that sustain effector function without succumbing to terminal exhaustion.

We next employed SCENIC analysis to highlight transcriptional regulatory systems that are most pertinent to the functional and prognostic characteristics of these T cell populations and revealed distinct regulatory patterns across the T cell subclusters, highlighting the differential activity of several key transcription factors (**FIGURE 4H <u>and SUPPLEMENTARY FIGURE 3D</u>**). *Jdp2*, an AP-1 repressor, exhibited differential regulon activity across the T cell subclusters, being elevated in subcluster 13 compared to subclusters 14-16. Higher *Jdp2* expression contributes to impaired cellular functionality by T cell exhaustion and anergy [40,41]. Therefore, lower activity of the *Jdp2* regulon in subclusters 14 and 15 is consistent with a more functional and active T cell state and, considering the enrichment of these subclusters with Combination treatment supports the notion that decreased *Jdp2* activity facilitates better immune responses. Furthermore, elevated activity of the *Mxd1* regulon in subcluster 13 suggests this, too, has a crucial role in maintaining the exhausted state characteristic of this subcluster by repressing cellular growth, contributing to its adverse clinical outcomes [42]. Antagonistic roles of *Mxd1* and *Myc* have been widely reported [43]. It is consistent, therefore, that the *Myc* regulon was notably active in T cell subclusters 14 and 15, whose signatures are linked to better prognoses. Elevated *Myc* activity in these clusters indicates a role that supports T cell persistence and functionality, perhaps through metabolic reprogramming and, when appropriate, proliferative activities that promote favorable immune responses.

The findings from our cytotoxic T cell subclustering and gene expression analyses are further supported by interrogation of bulk RNAseq data obtained from the YUMM2.1 subcutaneous tumors described earlier. From our previous study, tumors from IFNγ pre-treated cells are ‘immune deserts’ and, in accordance, were characterized by a marked reduction in effector T cell-associated signatures. In contrast, IgG-treated naïve tumors exhibited higher levels of T cell activation and exhaustion, reflecting an active yet ultimately ineffective immune response (**<u>SUPPLEMENTARY FIGURE 3F</u>**). Interestingly, tumors relapsing on α-PD-1 therapy displayed even higher levels of T cell activation and exhaustion than those treated with IgG alone, which suggests that despite the robust initial immune response, these tumors managed to evade sustained immune control. Conversely, α-PD-1 responding tumors displayed lower levels of exhaustion and elevated markers of T cell hyperactivation, consistent with a robust anti-tumor response. These bulk RNAseq findings align with our scRNAseq data, indicating that Combination therapy may successfully replicate the T-cell activation patterns seen in α-PD-1 responders. As the literature suggests activities that both elicit pro-inflammatory responses and mediate tumor progression for hyperactivated T cells [44,45], our TIDE analysis suggests the correlation with positive responses to α-PD-1 may be transient and insufficient for long-term tumor control, emphasizing the need for more durable therapeutic interventions.

To further explore the immune landscape shaped by Combination therapy, we performed a flow cytometry analysis of α-PD-1 relapsing tumors treated with either α-PD-1 or Combination therapy. Building on our observations of T cell dynamics, this revealed shifts in T cell populations following Combination treatment compared to α-PD-1 treated tumors (**<u>SUPPLEMENTARY FIGURE 3G</u>**). Specifically, Combination treatment reduced expression of GZMB by CD8^+^ T cells, suggestive of a larger proportion of this cell type transitioned to the quiescent state, as described previously in our scRNA-seq data. This shift was accompanied by nuanced changes in the expression of immune checkpoint markers. We observed a decrease in LAG-3 positivity among PD-1^+^ CD8^+^ T cells, indicative of reduced T cell exhaustion and an enhancement of immune response. However, this was counterbalanced by increased TIM-3 expression, another marker linked to T-cell dysfunction and exhaustion [46,47]. Together, these findings highlight the complexity and heterogeneity of T cell states in the TME that influence therapeutic efficacy.

### Combined α-PD-1 and PARP14 inhibitor treatment provokes dynamic immune evasion strategies in tumor cells

To further understand the impact of PARP14 inhibition on the TME, we further subclustered tumor cells and Cancer-Associated Fibroblasts (CAFs)— covering clusters 26-28— into six distinct subclusters (**FIGURE 5 A <u>and SUPPLEMENTARY TABLE 12</u>**), representing key roles in tumor progression and therapy adaptation. Among these, subcluster 3, identified as differentiation-primed tumor cells, was markedly enriched by α-PD-1 monotherapy, with its representation nearly doubling relative to IgG-treated tumors (**FIGURE 5B and C**). Slingshot trajectory analysis [14] positioned subcluster 3 as the likely origin of hyper-proliferative (subcluster 4) and immunomodulatory (subcluster 5) tumor states (**FIGURE 5D**). Gene Set Enrichment Analysis (GSEA) confirmed an abundance of differentiation-linked pathways in subcluster 3, suggesting that this state enhances susceptibility to immune-mediated elimination (**FIGURE 5E**). Notably, the suppression of this cluster by combination therapy underscores how targeting differentiation-associated vulnerabilities can limit tumor self-renewal and survival pathways.

**Figure 5.**
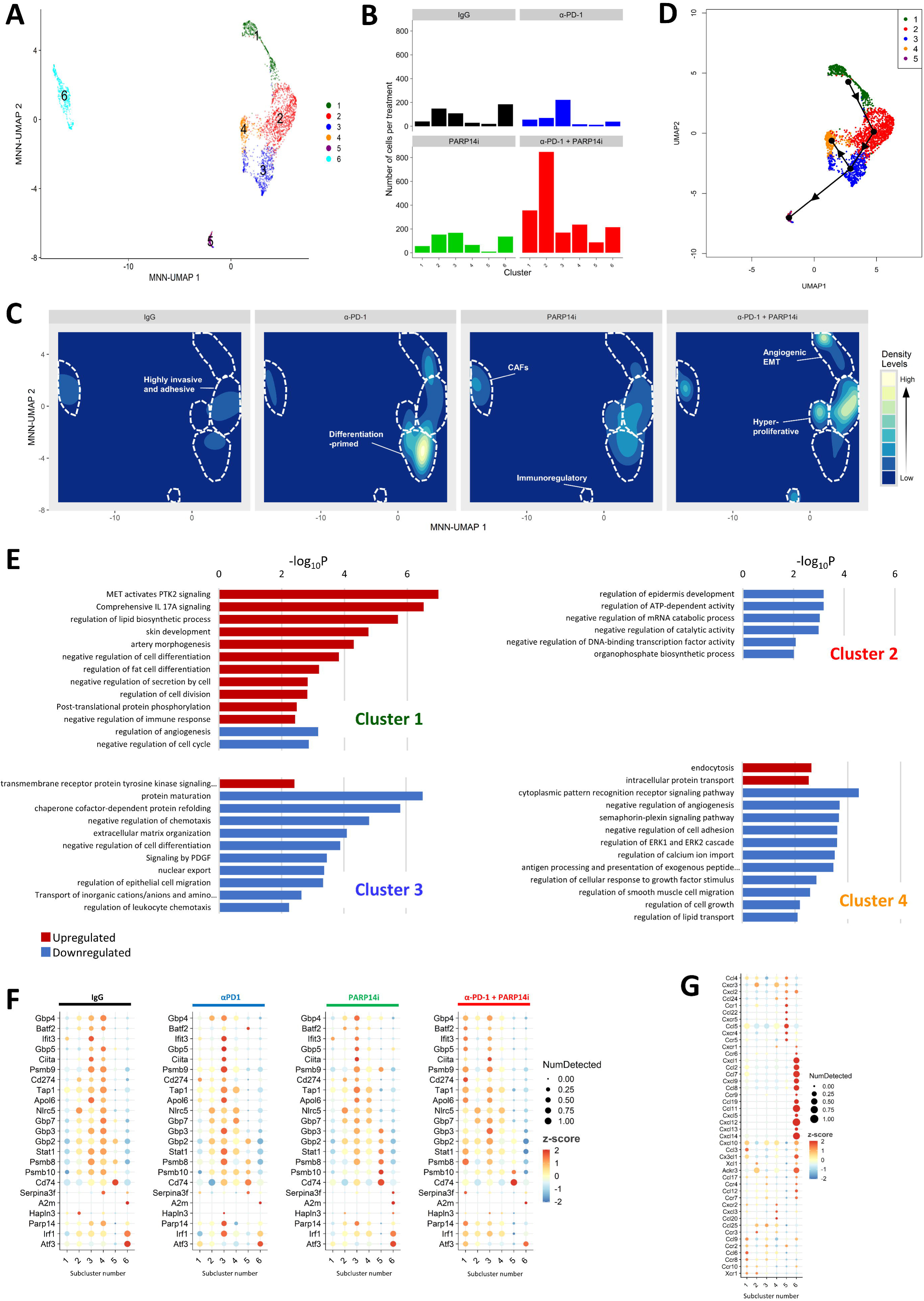
Combined treatment of α-PD-1 with PARP14 inhibitor promotes dynamic adaptive resistance strategies in tumor cells which permit their persistence. **A.** MNN corrected UMAP of tumor and cancer associated fibroblasts (CAFs) cells (clusters 26-28, *n=3,445*). Subclustered using Louvain method of community ordering and colored by subclusters. **B.** Cell number changes in each subcluster by treatment: IgG2a; α-PD-1; PARP14i (RBN012759); combination treatment α-PD-1 + PARP14i (RBN012759). **C.** Cell density projections of tumor and CAF cells by treatment. **D.** Slingshot trajectory inference analysis of tumor cells with trajectory direction indicated by arrows (black). **E.** Tumor subcluster identity, GSEA relevant terms of up-(red, FDR=0.05, logFC > 1) and downregulated (blue, FDR=0.05, logFC < −1) genes in tumor subclusters 1-4 (Metascape analysis, values presented as -log_10_ of P value). **F.** PARP14i signature gene expression presented as high (red) to low (blue) average expression and ratio of cells expressing (circle size), across tumor and CAF subclusters in different treatments. **G.** Cytokine and Chemokine gene expression presented as high (red) to low (blue) average expression and ratio of cells expressing (circle size), across Tumor and CAF subclusters.

In contrast, tumor subclusters 1, 2, and 4 were enriched by Combination treatment. Subcluster 2, characterized by highly invasive and adhesive tumor cells, drives metastasis via extracellular matrix remodeling and, we infer, an adaptive response to evade immune attack under intense therapeutic pressure, as evidenced by increased activation of biosynthetic pathways (**FIGURE 5E**). Subcluster 1, defined by angiogenic and epithelial-mesenchymal transition (EMT) phenotypes, enhances tumor migration, vascularization, and metastasis while also dampening immune responses (**FIGURE 5E**). The metabolic activation observed in subcluster 4 indicates an enhanced proliferative potential and cellular turnover under treatment-induced stress, consistent with therapy-adapted, resistant tumor states. These observations reveal distinct adaptive responses, including elevated angiogenesis, cell expansion, and interference with immune cell recognition machinery (**FIGURE 5E**). Therefore, while treatment suppresses key tumor-promoting pathways, it can also select tumor adaptations that sustain tumor growth and permit immune escape despite the pressure of combination therapy.

Integrative analysis of the top 200 DEGs across tumor subclusters allowed us to extract those uniquely regulated in each subcluster, highlighting distinct transcriptional patterns associated with therapeutic outcomes. By evaluating their expression in our YUMM2.1 bulk tumor RNAseq data set, we observed that genes enriched in subclusters 1, 2, and 5 correlated with relapse following α-PD-1 monotherapy, whereas those from subclusters 3 and 4 were linked to initial therapeutic responses (**<u>SUPPLEMENTARY FIGURE 5A</u>**).

Analysis of these unique gene lists using TIDE largely corroborated our identification of relapse or response outcomes. Subcluster 4, strongly associated with responders to α-PD-1 treatment in our data, was associated with improved patient survival (**<u>SUPPLEMENTARY FIGURE 5B and SUPPLEMENTARY TABLE 13</u>**). In contrast, gene signatures linked to subclusters relapsing on α-PD-1 therapy were predictive of worse clinical outcomes. Interestingly, genes uniquely upregulated in subcluster 3 did not reliably predict clinical response. This suggests that cells within this subcluster can adapt under selective pressure, transitioning into states that are either vulnerable to immune attack or capable of immune evasion. The dynamic interplay between these states ultimately shapes the overall treatment outcome.

Evaluation of the PARP14i gene signature across subclusters revealed treatment- and subcluster-specific regulation of key genes (**FIGURE 5F**). Upregulation of many of these in subcluster 1 by Combination therapy indicated elevated interferon signaling, while in subcluster 4, their downregulation suggests reduced metabolic and proliferative activities. In subcluster 3, α-PD-1 alone induced the PARP14i signature, but Combination therapy tempered its upregulation, possibly to prevent chronic stimulation. In subcluster 2, PARP14 inhibition selectively reduced *Ifit3* while increasing *Gbp7,* indicating a shift towards Type II interferon signaling.

## Discussion

Current melanoma therapeutics, particularly immune checkpoint inhibitors (ICIs) such as α-PD-1, have significantly advanced treatment, yet continue to face challenges from tumor resistance. Although combination therapies and targeted agents have been explored, their success in overcoming resistance mechanisms has been limited and necessitates novel treatment intervention approaches to enhance therapeutic efficacy [48,49]. Our previous research demonstrated PARP14 as a crucial regulator of immune responses, demonstrating that its inhibition amplifies the activity of infiltrating T cells within the TME [5]. In this study, we explore the effects of combining PARP14 inhibition with α-PD-1 therapy, to uncover the role of PARP14 upon immune cell dynamics in the tumor immune infiltrate at the onset of α-PD-1 resistance.

T cell exhaustion, marked by upregulation of inhibitory receptors and a reduction in cytokine production and cytotoxic activity, is a central challenge in overcoming resistance [50,51]. Our findings reveal that combining α-PD-1 with PARP14 inhibition mitigates T cell exhaustion by inducing a hyperactivate T cell state which can progress to quiescence. Elevated expression of genes associated with T cell hyperactivation contributes to a pro-inflammatory environment which can control tumor growth [45], yet if excessive, instead exacerbates tumor progression [44]. This implies an exit strategy, such as entering quiescence, preserves T cell functionality, preventing terminal exhaustion while maintaining a reservoir of reactivatable cells. These quiescent T cells, expressing longevity markers like *Bcl2* and *Il-6*, contrast with classical exhaustion and may sustain anti-tumor immunity. Leveraging this quiescent state could lead to durable responses by reactivating T cells with additional therapies.

Contact between PILRA and CD8α has been shown to maintain cytotoxic T cells in a quiescent state [52], presenting a potential axis for intervention. Moreover, *PILRA* expression correlates with tumor immune infiltration and positive prognosis [53], suggesting its role in preventing T cell exhaustion by promoting quiescence. Intriguingly, *Pilra* expression was enhanced in M1-like macrophage subclusters (**FIGURE 3F**) enriched by Combination treatment, supporting its role in inducing T cell quiescence. Notably, CD8α was highly expressed in T cell subclusters 14 and 16 (**<u>SUPPLEMENTARY FIGURE 3A</u>**), suggesting that overcoming PILRA inhibition may be required to exit the quiescent state. While monoclonal antibodies targeting PILRA have shown promise in preclinical models of autoinflammatory diseases [54,55], their potential in cancer treatment remains unexplored. Targeting the PILRA-CD8α axis therefore represents an appealing opportunity to enhance T cell-mediated immunity when combined with existing therapies, promoting more active anti-tumor responses.

In keeping with previous findings [7,8], we observed that PARP14 inhibition reshaped the TME by reducing immunosuppressive signals, typically linked with TAMs, and fostering a pro-inflammatory environment characterized by increased M1 macrophage polarization. TAMs support tumor progression through tissue repair and immunosuppression [56,57], while M1 macrophages are instrumental in sustaining effective anti-tumor immune responses via their production of pro-inflammatory cytokines such as TNF-α, IL-1β, and IL-6 [58,59]. Enriching M1 macrophages with PARP14i boosts cytokine and chemokine production, enhancing the effectiveness of PD-1 blockade therapy by increasing the infiltration and activation of cytotoxic T cells within the tumor [60,61]. Thus by enriching M1 macrophages, PARP14 inhibition can amplify T cell activation and improve α-PD-1 therapy, offering a potential strategy to overcome tumor immune evasion [62]. Together, the effects of PARP14 inhibition upon macrophages also likely contribute to counteracting T cell dysfunction often associated with markers of ICI resistance, such as chronic IFNγ exposure.

PARP14 influences gene expression and chromatin remodeling via its interactions and regulation of key immune pathways such as *STAT1* [8], *STAT3* [63], and *STAT6* [64] in co-operation with histone deacetylases (HDACs). Recent studies highlight the critical role of epigenetic regulation for macrophage polarization, whilst quiescent T cells actively manage their epigenetic landscape to preserve their long-term functionality whilst remaining poised for rapid reactivation [65]. We speculate that PARP14 inhibition disrupts the recruitment of HDACs to these regions, thereby promoting a “dormant” memory state in both T cells and macrophages. This state therefore requires further intervention to unlock the anti-tumor potential of these immune cell states.

Our study revealed a robust PARP14 inhibition (PARP14i) gene signature that is strongly associated with IFNγ signaling and STAT/IRF pathway regulation. While upregulated in α-PD-1-treated samples, this signature did not differentiate treatment response from relapse in our preclinical model. Nonetheless, high expression of the genes induced by PARP14i correlates with improved patient survival. While combination therapy enhances T cell activation and inflammation, it also induces expression changes amongst the PARP14i gene signature, which, combined with mounting immune pressure, may simultaneously facilitate the development of resistance mechanisms to allow tumor persistence. Notably, the upregulation of *Stat1*, *Atf3*, *Irf1*, and *Cd274* in subcluster 1 suggests that combination treatment may inadvertently enhance EMT-associated properties, thereby promoting more invasive and immune-evasive tumor cells [66]. These highly dynamic adaptive resistance mechanisms, therefore limit the promising effects our novel therapeutic combination upon immune cells. This may explain why despite short-term control of tumor growth in Combination therapy, these mechanisms later allow them to evade immunotherapy and progress despite intensified immune pressure. Conversely, the downregulation of IFNγ-regulated genes within the PARP14i signature in tumor subcluster 4 suggests reduced intrinsic IFN signaling. This reduction could heighten the susceptibility of these highly proliferative tumor cells to immune-mediated destruction [67]. This observation highlights the remarkable adaptability within the TME and aligns with findings from other studies that have reported similar resistance-associated tumor dynamics in response to targeted therapies [68,69].

The observed adaptive resistance underscores the necessity for a multifaceted approach to cancer treatment. While combining α-PD-1 therapy with PARP14 inhibition initially shows promise, rapid evolution of adaptive resistance mechanisms remains a considerable concern. To address this challenge, future research should focus on multi-modal strategies that integrate immune checkpoint inhibitors with agents that enhance pro-inflammatory immune infiltrates to produce a robust and durable anti-tumor response. Moreover, our identification of a PARP14i gene signature offers therapeutic promise, particularly for patients with low expression of these genes, enabling a more personalized treatment strategy. By using this signature as a biomarker, therapy can be tailored to individual patients in order to optimize treatment outcomes and minimize unnecessary interventions in patients who are less likely to respond. Further refinement should focus upon identifying complementary interventions to apply alongside the combination therapy that prevents tumor cells from evading immune detection or restores T cell-APC interactions to increase their tumor killing capacity. With continued research into resistance mechanisms and alternative pathways, robust combination therapies can potentially be developed which overcome even the most resilient tumor populations, offering a path toward more durable, comprehensive responses and improved patient survival.

## Declarations

## Acknowledgments

This study was funded by a research sponsorship agreement between Ribon Therapeutics and The University of Manchester, grant 821034 from the Melanoma Research Alliance (MRA), grant 22-0091 from Worldwide Cancer Research (WCR), and an award from the Skin Cancer Research Fund (SCaRF). C.W.W. was funded by a scholarship from the Hong Kong Scholarship for Excellence Scheme (HKSES). We thank Dr Gareth Howell and The University of Manchester Flow Cytometry Core Facility for facilitating flow cytometry analysis, Dr Andy Hayes and Claire Morrisroe in the Genomic Technologies Core Facility for assistance with single-cell RNA sequencing, Dr Leo Zeef and the Bioinformatics Core facility for their help with bioinformatic analysis, and members of the Biological Services Facility at The University of Manchester for help with animal work. FIGURE 2A and Graphical Abstract were created using BioRender.

## Author Contributions Statement

A.H. conceived the study. R.L., K.N.S., and C.W.W. performed experiments. I.L. and R.L. analyzed the scRNAseq data. K.N.S. analyzed the flow cytometry and bulk RNAseq data. D.T.I., M.N., and A.H. contributed to experimental design and data analysis. R.L., K.N.S., and A.H. drafted the original version of this manuscript, with all authors reviewing subsequent drafts. A.H. obtained funding for the study.

## Competing Interests Statement

The authors declare the following competing interests: D.T.I. and M.N. were employees and shareholders of Ribon Therapeutics at the time of data collection. A.H. received research sponsorship from Ribon Therapeutics. All other authors declare no competing interests.

## Supplementary Figure Legends

**Supplementary Figure 1 – Identification of a genetic signature associated with PARP14 inhibition.**

**A.** MC38 or A375 cells treated for 48h *in vitro* with either DMSO, 100nM PARP14 inhibitor (RBN012759), or 500nM PARP14 PROTAC (RBN012811); *n* = 3. Bulk RNAseq analysis of a PARP14 inhibition signature identified across human and mouse cell lines. Heatmap of treatment vs control per cell line, colors representing row-scaled z-score expression high (red) to low (blue). **B.** Clustergram of gene-TF associations derived from EnrichR analysis using the ARCHS4_TFs_Coexp database. Heatmap displays associations between genes (rows) and transcription factor terms (columns). Color-coded to represent the -log10 transformed adjusted p-values of enrichment, darker colors indicating stronger statistical significance. No normalization is applied, ensuring raw association strengths are visualized.

**Supplementary Figure 2 - A.** YUMM2.1 cells were implanted subcutaneously into the flank of female C57BL/6 mice. Once tumors reached ∼80mm^3^ in volume, treatment of α-PD-1 antibodies was commenced for a total of two doses, 3 days apart (horizontal dotted line). Tumor shrinkage and regrowth was monitored and individual growth curves plotted. When tumors reached a volume between ∼100-150mm^3^, a second round of treatment commenced (day 0); consisting of four IgG2a, α-PD-1, PARP14i or a combination of α-PD-1 + PARP14i. Treatment lasted 7 days. Tumors were subsequently excised and processed for single-cell RNAseq. **B.** RNAseq data of excluded IgG2a control treated tumor sample RL11i (black triangles and high-density lines) projected on the sample UMAP (color) using projecTILs R package. **C.** Sample representation in each cluster – histogram and **D.** UMAP. **E.** Cell number changes in each cell type by treatment: three mice IgG2a; three mice α-PD-1; three mice PARP14i (RBN012759) alone; and four mice a combination of α-PD-1 + PARP14i (RBN012759).

**Supplementary Figure 3- A.** Selected T cell marker gene expression presented as high (red) to low (blue) average expression and ratio of cells expressing (circle size), across T cell subclusters. **B.** Bulk RNAseq analysis on naïve YUMM2.1 tumors treated with IgG or α-PD-1. α-PD-1-treated cohort was split into relapsed (REL) and responders (RES). Compared to YUMM2.1 tumors pre-treated with chronic IFN-γ stimulation. Heatmap representations of activation and exhaustion markers. **C.** Gene expression presented as high (red) to low (blue) average expression and ratio of cells expressing (circle size), across T cell subclusters 13-16. **D.** Binarized heatmap of CD8+ T cell subclusters in Jdp2, Myc and Mxd1 regulons by relative regulon activity (SCENIC analysis). **E.** KM curves showing cancer patients with high TIDE scores of genes from T cell subclusters 13-16 and their effect on OS rates in bladder cancer cohort. Red indicates patients with high (top 50%) TIDE scores and blue indicates low (bottom 50%) TIDE scores. **F.** Representative image of Arid3a, Cebpb, Irf5, Mxd1 regulon activity AUC score in MNN corrected T cell subcluster UMAP. Calculated by SCENIC. **G.** Flow cytometry analysis of YUMM2.1 tumor xenografts in C57BL/6 mice relapsing on an initial dose of α-PD-1 and harvested on-treatment following subsequent dosing with either α-PD-1 (Blue; *n* = 6) or α-PD-1 + PARP14i (Red; *n* = 6). **From left to right:** Percentage of total immune cells and individual T cell subsets as a proportion of CD45^+^ cells. Percentage of cytotoxic CD8^+^ T cells evaluated via their expression of GZMB. Percentage of exhaustion subsets in cytotoxic T cells measured by LAG-3 and TIM-3 levels for PD-1^+^, PD-1^+^CTLA-4^+^, or PD-1^+^CTLA-4^−^ background. The p-value was assessed by two-sided unpaired t-test and the data were presented as the meanllJ±llJSEM. \**p* < 0.05, \*\*\**p* < 0.01, \*\*\**p* < 0.001, \*\*\*\**p* < 0.0001.

**Supplementary Figure 4 – Flow cytometry gating strategy for T cell subsets and exhaustion phenotypes**

**Supplementary Figure 5 – A.** Bulk RNAseq analysis on naïve YUMM2.1 tumors treated with IgG or α-PD-1. α-PD-1-treated cohort was split into relapsed and responders. Compared to YUMM2.1 tumors pre-treated with chronic IFN-γ stimulation. Heatmap representations of uniquely upregulated genes per each subcluster within the top 200 expressed genes. Boxes represent relevant clinical outcome per cluster. **B.** KM curves showing cancer patients with high TIDE scores of uniquely upregulated genes per tumor subcluster and their effect on OS rates in bladder cancer cohort. Red indicates patients with high (top 50%) TIDE scores and blue indicates low (bottom 50%) TIDE scores.

## Supplementary Figure Legends

Supplementary Table 1: Gene Signatures

Supplementary Table 2: Tumor Size

Supplementary Table 3: Sample Identity

Supplementary Table 4: Cell Number Per Sample

Supplementary Table 5: Cluster Identity

Supplementary Table 6: Cell Number Per Cluster

Supplementary Table 7: Expression of Manually Selected Genes

Supplementary Table 8: Top 100 Expressed Genes Per Cluster – Main

Supplementary Table 9: Top 100 Expressed Genes Per Cluster - NK and T Cells

Supplementary Table 10: Top 25 Genes Up / Down Regulated in T Cell Clusters 13-16

Supplementary Table 11: Top 100 Expressed Genes Per Cluster - Mono / Macs

Supplementary Table 12: Top 100 Expressed Genes Per Cluster - Tumor and CAFs

Supplementary Table 13: Uniquely Expressed Genes Tumor

## Notes

### Summary of Updates

We have clarified how combination therapy enriches cytotoxic T-cell subclusters and shifts macrophage polarisation towards an M1-like state. We emphasise the necessity of further intervention to fully exploit these immune populations due to tumour adaptive resistance. We expanded our analysis on T-cell migration into the tumour microenvironment, detailing chemokine-receptor interactions driving immune cell recruitment and reprogramming. Particular attention was given to subclusters most responsive to combination therapy. To improve flow and readability, we restructured sections Results, Discussion.

