## Supplementary figures and images for "Combined PARP14 Inhibition and PD-1 Blockade Promotes Cytotoxic T Cell Quiescence and Modulates Macrophage Polarization in Relapsed Melanoma"

### Supplementary figures

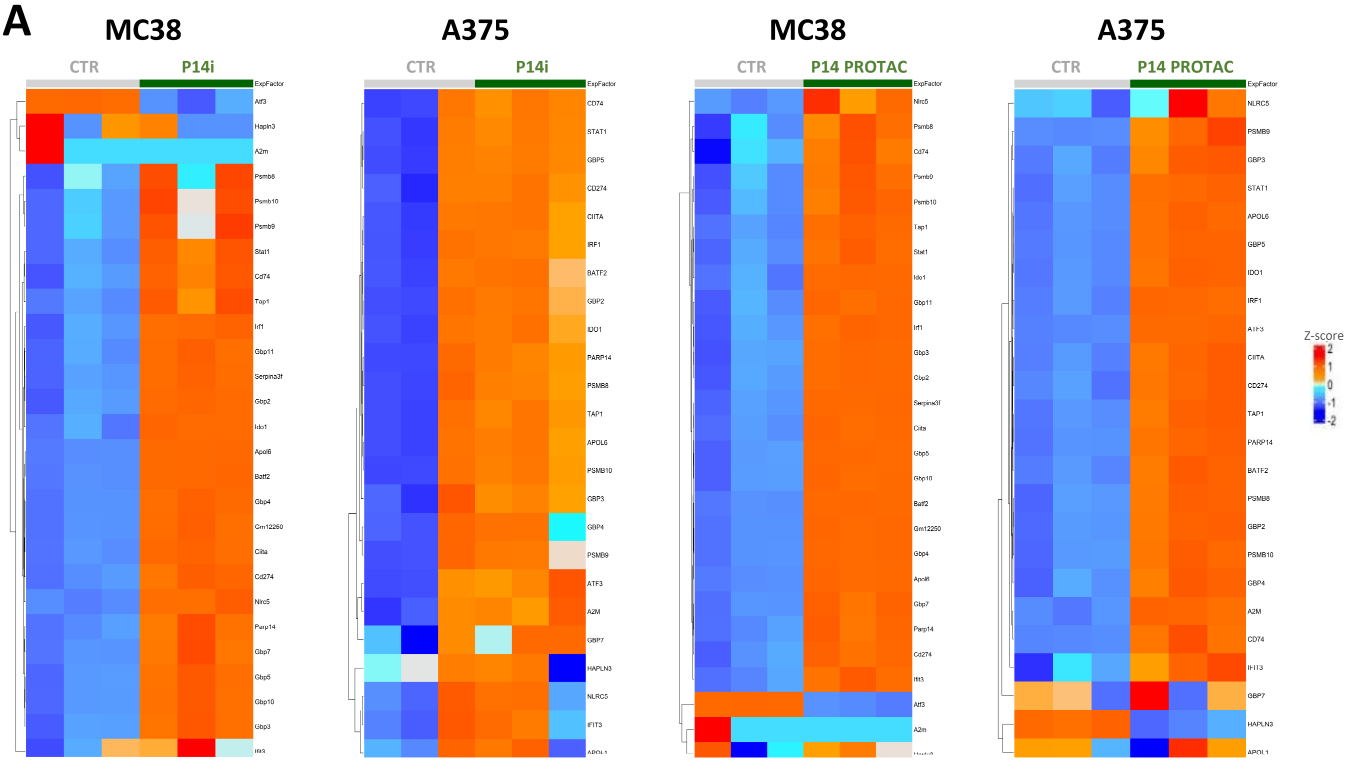

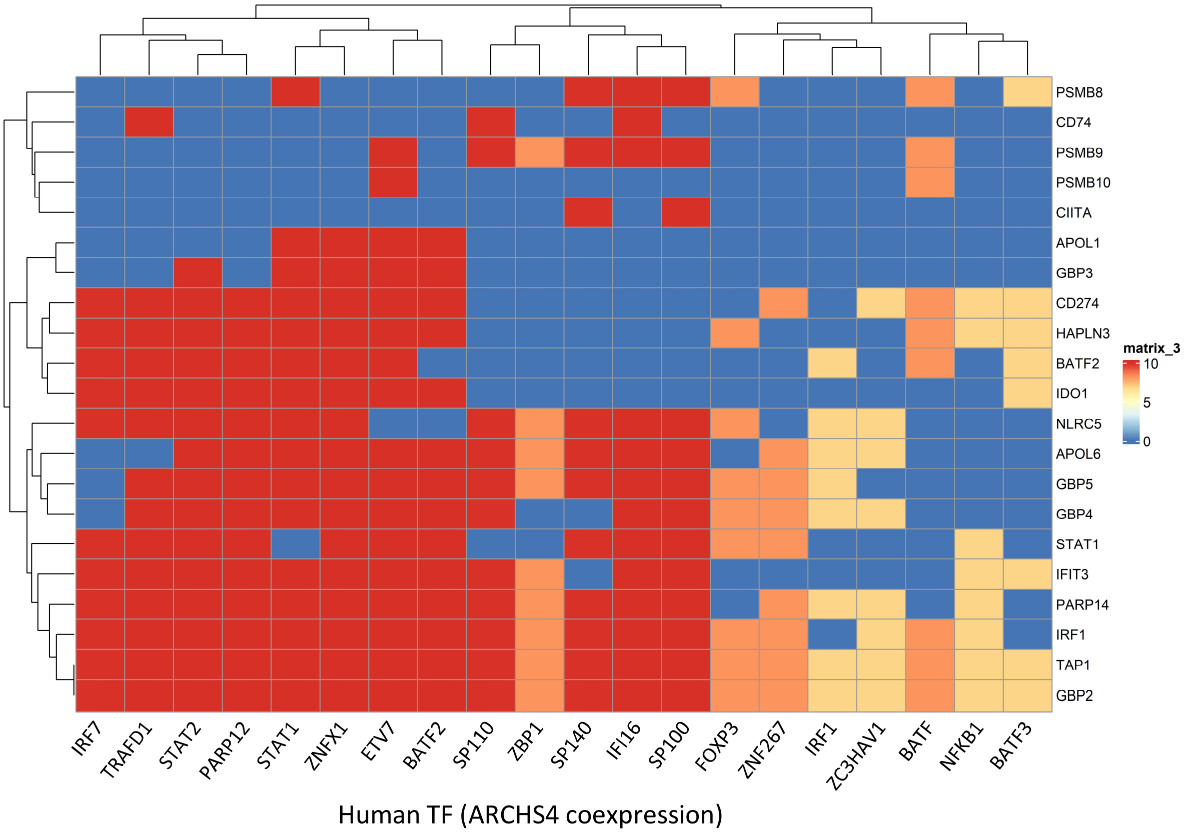

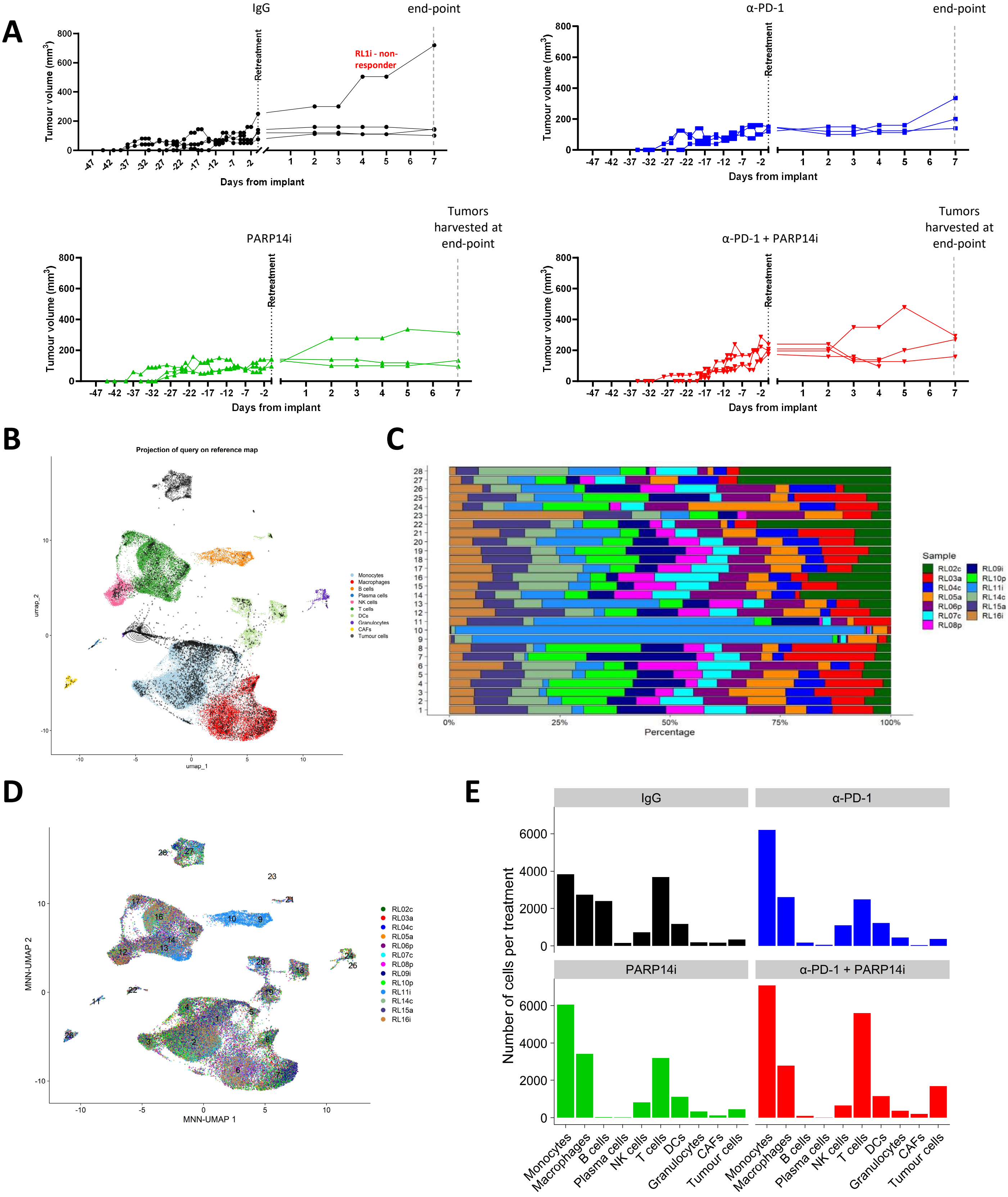

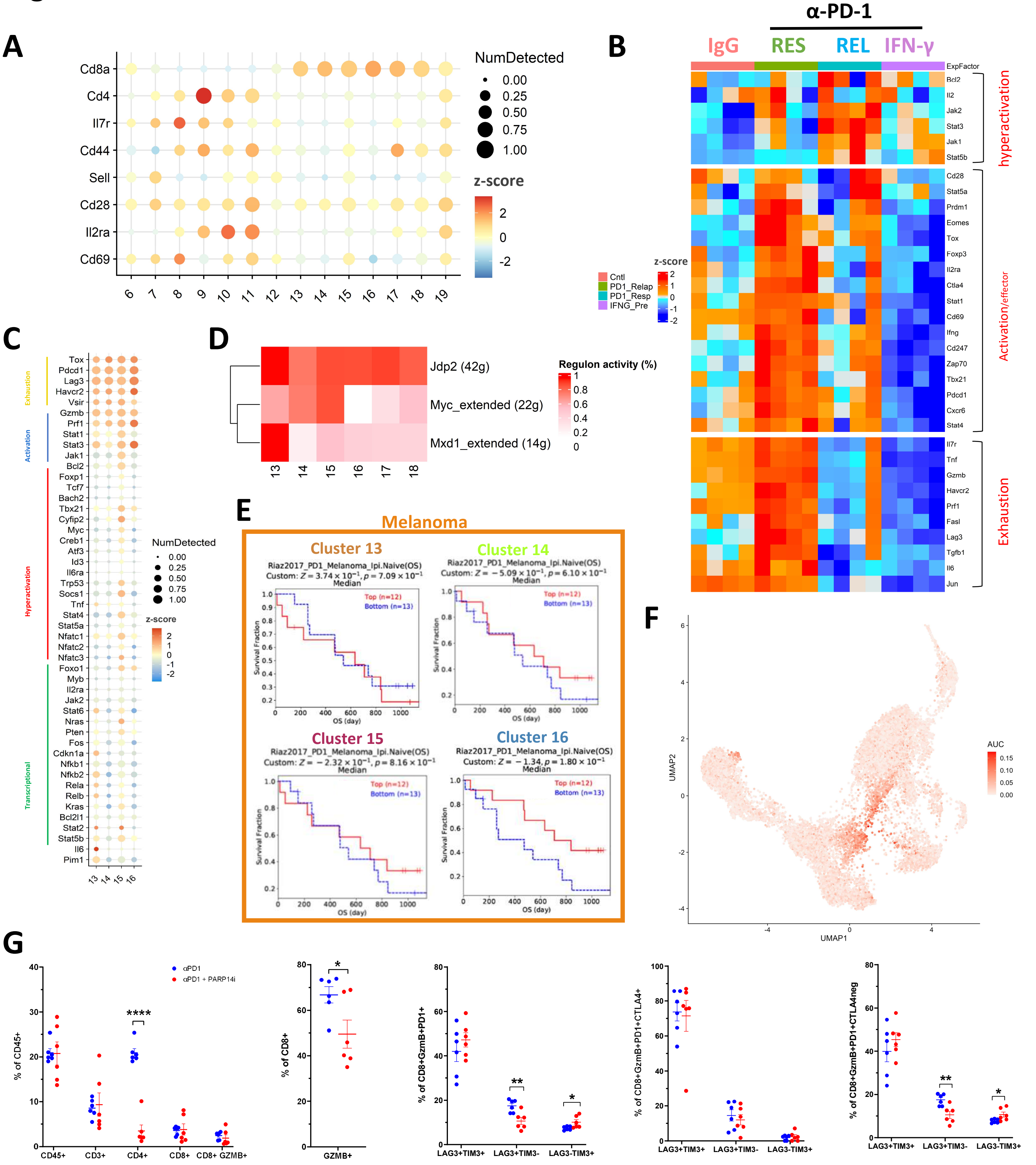

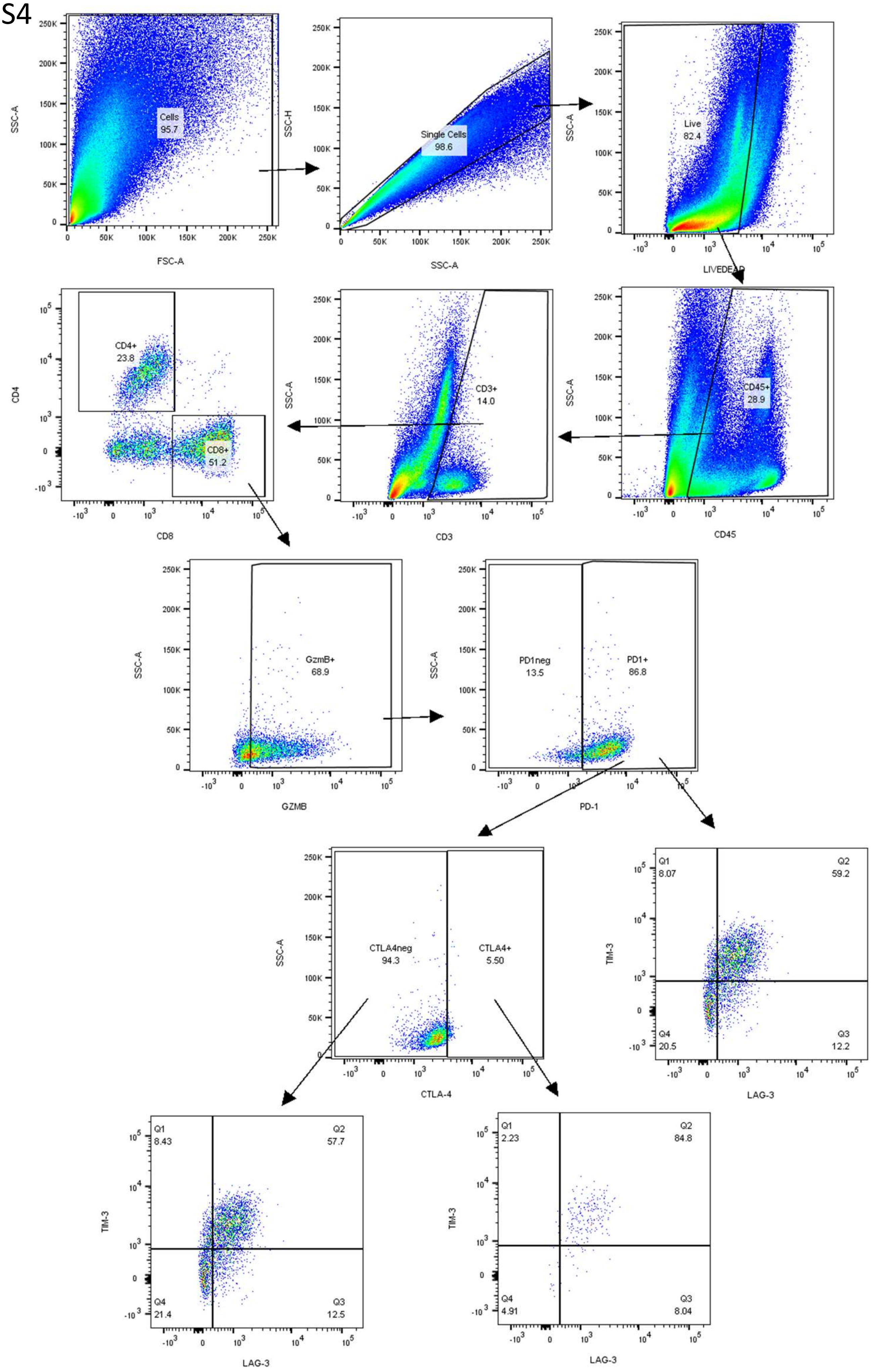

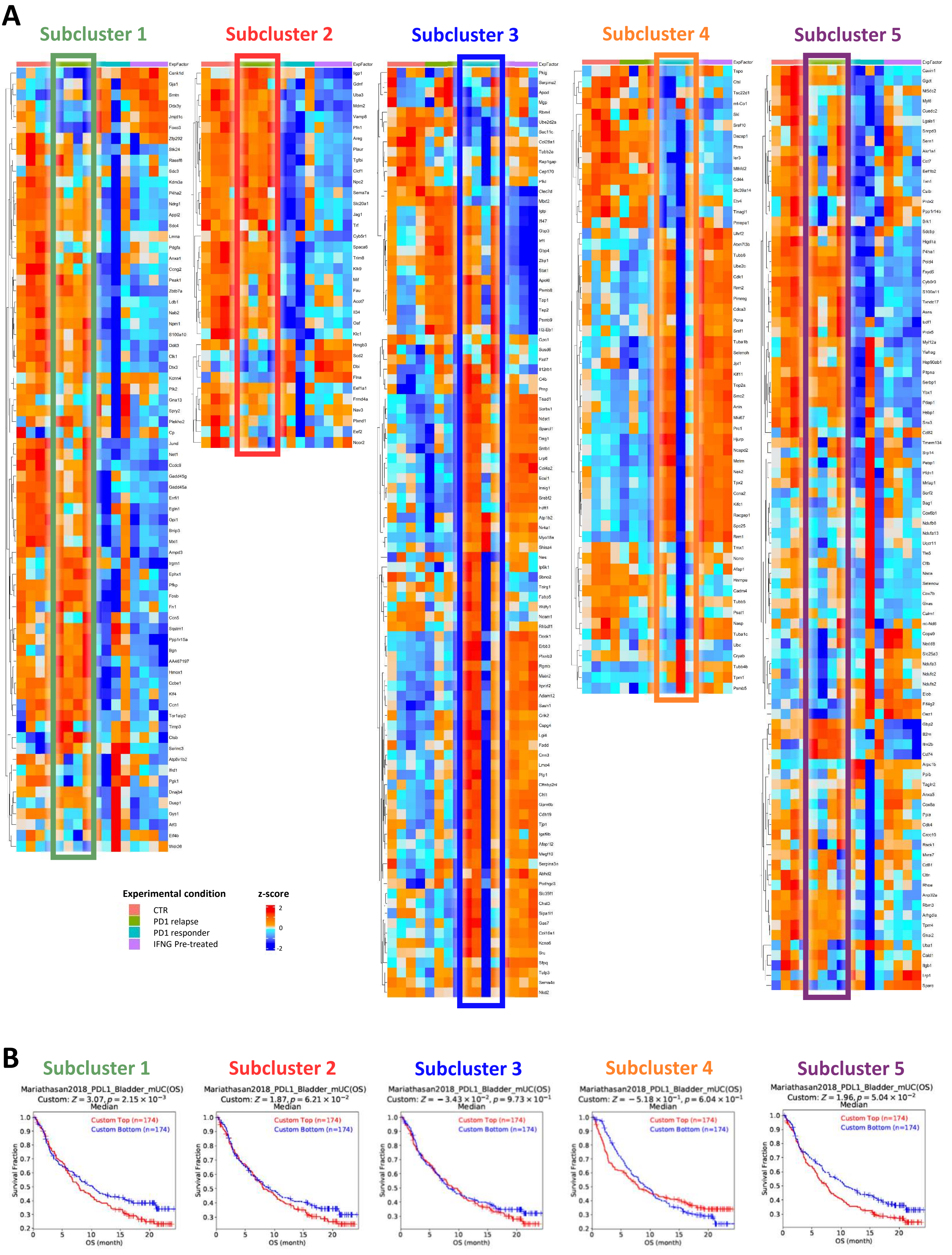
